# Basolateral amygdala parvalbumin neurons report aversive prediction error to constrain fear learning

**DOI:** 10.1101/2020.09.22.307561

**Authors:** Joanna Oi-Yue Yau, Chanchanok Chaichim, John M. Power, Gavan P. McNally

**Author notes:** Correspondence to: Gavan P. McNally PhD, School of Psychology, UNSW Sydney, NSW Australia.

## Abstract

Animals, including humans, use prediction error to guide learning about danger in the environment. The basolateral amygdala (BLA) is obligatory for this learning and BLA excitatory projection neurons are instructed by aversive prediction error to form fear associations. Complex networks of inhibitory interneurons, dominated by parvalbumin (PV) expressing GABAergic neurons, form the intrinsic microcircuitry of the BLA to control projection neuron activity. Whether BLA PV interneurons are also sensitive to prediction error and how they use this error to control fear learning remains unknown. We used PV cell-type specific recording and manipulation approaches in male transgenic PV-Cre rats to address these issues. We show that BLA PV neurons control fear learning about aversive events but not learning about their omission. Furthermore, during fear learning BLA PV neurons express the activity signatures of aversive prediction error: greater activity to unexpected than expected aversive events and greater activity to better rather than poorer predictors of these events. Crucially, we show that BLA PV neurons act to limit fear learning across these variations in prediction error. Together, this demonstrates that prediction error instructs and regulates BLA fear association formation in a cell-type specific manner. Whereas BLA projection neurons use prediction error signals to form and store fear associations, BLA PV interneurons use prediction error signals to constrain fear association formation.

**Significance Statement:** The capacity to predict sources of danger in the environment is essential for survival. This capacity is supported by associative learning mechanisms that are triggered when the danger experienced is greater than the danger expected. Here we show that the activity of parvalbumin positive GABAergic interneurons in the rat basolateral amygdala neurons report this difference between the danger expected and the danger experienced and that they use this difference to limit the amount of fear which is learned.

---

Animals, including humans, use prediction error to guide learning about danger in the world (Fanselow, 1998; Fanselow and Poulos, 2005; Li and McNally, 2013; Wright et al., 2019; Yau and McNally, 2018a; Yau and McNally, 2018b). Prediction error determines how effective aversive events are in supporting fear learning. An unexpected aversive unconditioned stimulus (US), such as a painful shock, produces high prediction error to promote fear learning whereas the same aversive event when expected produces low prediction error and does not support fear learning (Eippert et al., 2012; Kamin, 1968).

Basolateral amygdala glutamatergic projection (principal) neurons are obligatory for this fear learning and their activity during fear learning reports the difference between the expected US and the US actually received, the aversive prediction error (Herry and Johansen, 2014; McNally et al., 2011). In rats, single unit recordings show greater activity to unexpected than expected aversive USs (Johansen et al., 2010; Ozawa et al., 2017). Similar findings have been reported using other measures of BLA neuronal activity in rats (Furlong et al., 2010), calcium transients in loaded principal neurons in mice (Krabbe et al., 2019) as well as tissue oxygenation in mice (McHugh et al., 2014) and humans (Dunsmoor et al., 2007, 2008; Eippert et al., 2012; Michely et al., 2020). These reductions in aversive US-related activity are accompanied by increases in activity to the fear conditioned stimulus (CS) predicting expected aversive events (Eippert et al., 2012; Johansen et al., 2010; Michely et al., 2020). Importantly, variations in US-evoked BLA projection neuron activity are causal to variations in fear learning because restoring their activity to an expected shock US also restores fear learning about that shock US (Sengupta et al., 2016). Nonetheless, how BLA projection neurons achieve this sensitivity to prediction error remains poorly understood.

The BLA is comprised of complex, intrinsic microcircuitries of inhibitory interneurons. These interneurons are distinct families of GABAergic neurons defined by expression of a variety of markers (e.g., somatostatin, cholecystokinin, and vasoactive-intestinal peptide (VIP)) of which parvalbumin (PV) containing interneurons are the most common (McDonald, 1992; McDonald and Betette, 2001; Sah et al., 2003). BLA PV interneurons serve a key role in fear learning. They adaptively and dynamically gate projection neuron activity, synaptic plasticity, and fear learning (Krabbe et al., 2018; Krabbe et al., 2019; Letzkus et al., 2015; Lucas et al., 2016; Polepalli et al., 2020; Polepalli et al., 2010; Wolff et al., 2014; Woodruff and Sah, 2007a, b). In other brain regions, the actions of local GABAergic neurons are essential for computing prediction error. For example, reward prediction error computation in the ventral tegmental area (VTA) depends on the activity of GABAergic interneurons. These interneurons provide local synaptic inhibition that controls and shapes VTA dopamine neuron responses to the rewarding US and the learning instructed by these responses (Cohen et al., 2012; Eshel et al., 2015). Whether BLA PV GABAergic interneurons serve a similar role in computing aversive prediction error and instructing fear learning by BLA projection neurons is poorly understood. There is evidence that aversive US-evoked activity in BLA VIP interneurons co-varies with US surprisingness, being smaller to expected than unexpected USs (Krabbe et al., 2019). But, how the activity of other BLA GABAergic interneurons co-vary with US expectancy and whether any BLA interneuron population has a causal role in implementing aversive prediction errors are unknown.

Here we used the recently developed PV-Cre rat (Wright et al., in preparation) to address these issues and determine the role of BLA PV neurons in aversive prediction error. We show that BLA PV neurons have dynamic and opposing changes in US- and CS-evoked activity across fear learning and that this activity of BLA PV neurons during fear conditioning reports an aversive prediction error. However, in contrast to other BLA neuron populations which use prediction error to form and store fear associations, we show that BLA PV neurons use this prediction error to constrain fear association formation.

## Materials and Methods

### Subjects

Male PV-Cre (LE-Tg(Pvalb-iCre)2Ottc) heterozygous rats (Optogenetics and Transgenic Technology Core, National Institute on Drug Abuse [NIDA], National Institutes of Health, MD, USA) or their Cre-littermates obtained from the NIDA IRP Transgenic Rat Project (Wright et al., in preparation) via the Rat Resource and Research Center (RRRC# 00773, Missouri, USA) and bred at the Animal Resources Centre (Perth, Australia). They were housed in groups of 2 to 4 in a colony room maintained on a 12:12 light dark cycle (lights on 7:00). Rats were maintained on 15g of standard lab chow per day with free access to water. Experiments were approved by the UNSW Animal Care and Ethics Committee and performed in accordance with the Animal Research Act 1985 (NSW), under the guidelines of the National Health and Medical Research Council Code for the Care and Use of Animals for Scientific Purposes in Australia (2013).

### Apparatus

Behavioral testing was conducted in Med-Associates chambers [24 cm (length) x 30 cm (width) x 21 cm height)] enclosed in ventilated, sound-attenuating cabinets [59.5 cm (length) x 59 cm (width) x 48cm (height)]. The left sidewall was fitted with a magazine dish where grain pellets (Bio-Serv, USA) were delivered when a lever located 4 cm to the right of the magazine was pressed. A 3 W houselight was mounted on top of the right wall provided illumination in the chamber throughout every session. An LED light mounted to the roof of the sound-attenuating cabinet was used to deliver a flashing visual CS. A speaker attached to the right-side wall of the chamber was used to deliver auditory CSs. A metal grid was fitted to the floor of the chamber to deliver the scrambled footshock US. For optogenetic experiments, an LED driver with an integrated rotary joint (Doric Instruments) was suspended above the center of the operant chamber.

### Surgeries and viral vectors

Rats were anaesthetized via 1.3ml/kg ketamine (Ketamil, Ilium) (100 mg/ml) and 0.3 mg/kg xylazine (Xylazil, Ilium) (20 mg/ml.) They received s.c carprofen (Rimadyl, Zoetis) and 0.5% bupivacaine under the incision site. A 5 μl, 30-gauge conical tipped microinfusion syringe (SGE Analytical Science) was used to infuse 0.75 μl of AAV vectors into BLA (A-P - 3.00; M-L ± 5.00; D-V -8.15 in mm from bregma) (Paxinos & Watson, 2007) at a rate of 0.25 μl/minute (UMP3 with SYS4 Micro-controller, World Precision Instruments) and the syringe was left in place for an additional 7 minutes. At each craniotomy, a fiber optic, ceramic cannula (Thorlabs) was lowered into the BLA (A-P -3.00; M-L ± 5.10; D-V -8.00) (Paxinos & Watson, 2007) and secured in place by dental cement anchored to the screws and the skull. The incision was sutured and i.p. antibiotic (Duplocillin, Intervet) applied. Rats were monitored until the end of the experiment.

The Cre-dependent AAV vectors used were: AAV5-ef1-DIO-eYFP (3.3×10^12^ GC/ml) (pAAV-Ef1a-DIO EYFP was a gift from Karl Deisseroth (Addgene viral prep # 27056-AAV5; http://n2t.net/addgene:27056; RRID:Addgene_27056); AAV5-ef1α-DIO-hChR2(H134R)-eYFP (5.5×10^12^ GC/ml) (pAAV-EF1a-double floxed-hChR2(H134R)-EYFP-WPRE-HGHpA was a gift from Karl Deisseroth (Addgene viral prep # 20298-AAV5; http://n2t.net/addgene:20298; RRID:Addgene_20298); AAV5-ef1α-DIO-eNpHR3.0-eYFP (1.1×10^13^ GC/ml) (pAAV-Ef1a-DIO eNpHR 3.0-EYFP was a gift from Karl Deisseroth (Addgene viral prep # 26966-AAV5; http://n2t.net/addgene:26966; RRID:Addgene_26966); AAV9-Syn-FLEX-gCaMP7s (1.2×10^13^ GC/ml) (pGP-AAV-syn-FLEX-jGCaMP7s-WPRE was a gift from Douglas Kim & GENIE Project (Addgene viral prep # 104491-AAV9; http://n2t.net/addgene:104491; RRID:Addgene104491).

### Procedure

#### Anatomy

PV-Cre rats with Cre-dependent AAV vectors in the BLA (eNpHR3.0: n = 5; eYFP; n =3) were transcardially perfused with saline containing 1% sodium nitrate and heparin (5000 IU/ml) and paraformaldehyde (4%), pH 7.4. Brains were extracted, postfixed (1hr) and cryoprotected in 20% sucrose (48 h). Brains were frozen, and 40 μm BLA sections were sliced and collected on a cryostat (CM 1950, Leica) and stored in a 0.1 M PB saline solution containing 0.1% sodium azide at 4°C. Two color immunofluorescence was used to reveal PV as well as enhanced yellow fluorescent protein (eYFP) immunoreactivity (IR). Free-floating tissue was washed in PB, pH 7.4, blocked (2 h, 5% NGS in PBTX), and placed in 1:1500 chicken anti-GFP (Thermo Fisher Scientific Cat# A10262, RRID:AB_2534023) and 1:1000 mouse anti-PV (Sigma-Aldrich Cat# P3088, RRID:AB_477329), diluted in a solution of PBTX and 2% NGS at room temperature for 24 h. Sections were washed in PB for 20 min and then incubated in 1:1000 Alexa-488 goat anti chicken (Molecular Probes Cat# A-11039, RRID:AB_142924) and 1:750 Alexa-594 goat anti-mouse (Molecular Probes Cat# A-11032, RRID:AB_2534091) diluted in PBTX and 2% NGS at room temperature for 2 h. Sections were washed for 30 min in PB, mounted, and cover slipped with Permafluor (Thermofisher Scientific, MA, USA). The BLA of each rat was delineated according to Paxinos and Watson (2007) and across two sections, neurons positive for PV-IR and eYFP-IR assessed via Olympus BX53 upright microscope (Olympus, Shinjuku, Tokyo, Japan) and CellSens (Olympus) software and counted using Photoshop (Adobe).

### Electrophysiology

#### Slice preparation

Coronal brain slices (350 μm) were prepared from PV-Cre+ rats that received either AAV5-ef1α-DIO-eNpHR3.0-EYFP or AAV5-ef1α-DIO-hChR2(H134R)-eYFP to BLA. Rats were deeply anaesthetized with isoflurane (5%), decapitated and the brain removed and placed in ice-cold cutting ACSF (95 mM NaCl, 2.5 mM KCl, 30 mM NaHCO3, 1.2 mM NaH2PO4, 20 mM HEPES, 25 mM glucose, 5 mM ascorbate, 2 mM thiourea, 3 mM sodium pyruvate, 0.5 mM CaCl2, and 10 mM MgSO4). The brain was trimmed, sliced using a vibratome (model VT1200, Leica, Wetzlar, Germany), then incubated for 10-15 min at 30°C in the recovery ACSF (95 mM NMDG, 2.5 mM KCl, 30 mM NaHCO3, 1.2 mM NaH2PO4, 20 mM HEPES, 25 mM glucose, 5 mM ascorbate, 2 mM thiourea, 3 mM sodium pyruvate, 0.5 mM CaCl2, and 10 mM MgSO4), before being transferred a Braincubator (Payo Scientific, Sydney, Australia) and held at 18°C in holding-ACSF (identical to Cutting ACSF, but with 2 mM CaCl2, and 2 mM MgSO4). All solutions were pH adjusted to 7.3-7.4 with HCl or NaOH and gassed with carbogen (95% O2 - 5% CO2).

#### Whole-cell patch clamp recordings

Slices were transferred to a recording chamber and continuously perfused with standard ACSF (30°C) containing (in mM): NaCl, 124; KCl, 3; NaHCO3, 26; NaH2PO4, 1.2; glucose, 10; CaCl2, 2.5; and MgCl2, 1.3. Targeted whole-cell patch-clamp recordings were made from EYFP+ BLA neurons using a microscope (Zeiss Axio Examiner D1) equipped with 20x water immersion objective (1.0 NA), LED fluorescence illumination system (pE-2, CoolLED) and an EMCCD camera (iXon+, Andor Technology). Patch pipettes (3 – 5 MΩ) were filled with an internal solution containing 130 mM potassium gluconate, 0 mM KCl, 10 mM HEPES, 4 mM Mg2-ATP, 0.3 mM Na3-GTP, 0.3 mM EGTA, and 10 mM phosphocreatine disodium salt (pH 7.3 with KOH, 280 - 290 mOsm). Electrophysiological recordings were amplified using a Multiclamp amplifier (700B, Molecular Devices, California, USA), filtered at 6-10 kHz, and digitized at 20 kHz with a National Instruments multifunction I/O device (PCI-6221). Recordings were controlled and analyzed offline using Axograph (Axograph, Sydney, Australia). Liquid junction potentials were uncompensated.

Series resistance and membrane resistance were calculated using in built routines in Axograph. To determine the passive membrane, AP waveform and neuronal firing pattern, neurons were maintained at −65 mV using and a series of 600 ms current injections (−100 to 300 pA, 25 pA steps) was applied. The AP threshold was defined as the potential at which the AP velocity exceeded 10 mV / ms. The AP amplitude, half-width and fast afterhyperpolarization (fAHP) were calculated relative to threshold. eNpHR3.0 was stimulated using orange light (GYR LED bandpass filtered 605/50 nm) delivered through the objective. ChR2 was stimulated using blue light (470 nm LED). To determine the effect of photoinhibition on the AP firing frequency, neurons were induced to fire via depolarizing current injection, or trains (5 ms 10 Hz) of depolarizing current injections; the (amplitude of current pulse was set to 10 pA above the minimum current required to evoke an AP). When possible, protocols were repeated 5-10 times and the results averaged. Data were excluded if the series resistance was > 20 MΩ or more than 150 pA was required to maintain the neuron at −65 mV.

### Behavior

#### Baseline lever press training

All behavioral experiments used conditioned suppression as fear measure (Arico et al., 2017; Arico and McNally, 2014; Sengupta et al., 2016; Sengupta et al., 2018; Yau and McNally, 2015). Conditioned suppression has a non-zero baseline because rats lever press for a pellet reward at a constant rate and so can reveal decreases and increases in fear; there are high levels of baseline activity during training and testing sessions; it is equally sensitive to visual and auditory CSs despite these CSs eliciting different amounts of freezing (Bevins and Ayres, 1991); and assessment is fully automated. All experiments began with lever pressing training. On Day 1, rats received magazine training with lever presses rewarded on fixed ratio 1 [FR1] schedule in addition to free pellet rewards on a fixed interval (FI) 300 s schedule. Magazine training terminated at 60 min, or when the rat reached 100 lever presses. On Day 2, rats received FR1 lever press training. On Day 3, lever pressing was maintained on a variable interval (VI) 30 s schedule. From Day 4 until the end of the experiment, rats were maintained on a VI60 s. All sessions lasted 60 min unless otherwise noted. On Day 9 – 10, rats received CS pre-exposure. There were 4 presentations of each CS, with an ITI between 400 – 720 s. All rats were tethered to dummy patch cables on Day 7 – 9.

#### Fear acquisition

There were ChR2 (n = 7) or eNpHR3.0 (n = 8) groups. PV-Cre +/- rats transfected with eYFP (n = 4) were combined with a group of PV negative rats that received ChR2 (n = 5) to make group Control (n = 9). All rats underwent fear conditioning on Day 11 – 14. Rats were tethered to patch cables outputting at least 10mW of 625 nm or 465 nm light. Presentation of an auditory stimulus (85dB, 1Hz clicker) co-terminated with a 0.5s, 0.5mA footshock US. Delivery of 625 nm (for eNpHR3.0 and Control) and 465 nm light (for ChR2 and Control) flanked presentation of the US, beginning 0.5 s before shock onset and terminating 0.5 s after shock onset. There were 4 pairings per session, with a random intertrial interval (ITI) between 500 – 900 s. On Day 15 rats were tested for fear response to the CS. The CS was presented four times on a random ITI between 500 – 900 s. For intertrial interval inhibition, eNpHR3.0 (n = 7) or eYFP (n = 8) groups underwent fear conditioning as described above. Photoinhibition (1.5 s) occurred randomly during each ITI.

To determine the effects of pulsed versus continuous ChR2 excitation, PV-Cre +/- rats were spilt into two groups, ‘ChR2 constant’ and ‘ChR2 pulse’, whereas PV-Cre negative rats served as controls. All rats underwent fear conditioning to an auditory stimulus (85dB, 1Hz clicker) paired with a 0.5 s, 0.6mA footshock US on Day 11 – 13. Photostimulation flanked the US. ChR2 constant (n = 9) and Control (n =9) rats received stimulation as described above, whereas rats in the ChR2 pulse (n = 7) groups received a 25Hz pulsed stimulation, 50% duty cycle (20 ms on, 20 ms off) and rats in the Control group received either manipulation. Rats were tested for their fear response to the CS on Day 14. The CS was presented 4 times on an ITI between 500 – 900s.

#### Fear Extinction

eNpHR3.0 (n = 9) or eYFP (n = 7) groups underwent fear conditioning to a tone CS (85dB, 2800Hz) on Days 11 - 13, with presentations co-terminating with a 0.5 s, 0.6mA footshock. There were 4 pairings per day with an ITI of 500 – 900 s. Fear to the tone was then extinguished across four days with 4 presentations per day and an ITI of 500 – 900 s. Rats received continuous delivery of 625 nm light for 5 seconds at the time of US omission, beginning 0.5s prior to tone offset, and extending for an extra 4.5 s.

#### Fiber Photometry during blocking of Pavlovian fear

PV-Cre +/- expressing gCaMP7s were divided into two behavioral groups – Block (n=10) and Control (n = 9). Block groups received Stage I training on Days 11 – 13. Presentation of CSA (85dB 10Hz clicker) co-terminated with a 0.5 s, 0.6mA footshock. There were 4 presentations each day with an ITI between 600 – 720 s. The Control group received VI60 lever press training instead. All rats received Stage II training on Day 14 and 15. Prior to each session, rats were tethered to patch cables. In Stage II, CSA was simultaneously presented with CSB (1Hz flashing LED) and this compound cue was co-terminated with a 0.5 s, 0.6mA footshock. There were 4 presentations each day with an ITI between 600 – 720 s. On Day 16, rats were tested for their fear response to CSB. CSB was presented four times, with an ITI of 900 s in a 70 min session. Rats were tethered to dummy patch cables on Day 7 – 9.

Recordings were made during Stage II using Fiber Photometry Systems from Doric Lenses and Tucker Davis Technologies (RZ5P, Synapse). 465 nm (Ca2+ - dependent signal) and 405 nm (isosbestic control signal) emitted from LEDs controlled via dual channel programmable LED drivers (were channelled into 0.39 NA, Ø400μm core multimode pre-bleached patch cables. Light intensity at the tip of the patch was maintained at 10-30μW across sessions. GCaMP7 and isosbestic fluorescence were relayed via patch cables, amplified, and measured by Doric Fluorescence Detectors. Synapse software controlled and modulated excitation lights (465nm: 209Hz; 405nm: 331Hz), as well as de-modulated and low-pass filtered (3Hz) transduced fluorescence signals in real-time via the RZ5P. RZ5P/Synapse also received Med-PC signals to record behavioral events in real time.

#### Blocking of Pavlovian fear

eNpHR3.0 (n = 15) and eYFP (n = 15) groups were divided into two behavioral groups – Block (eYFP n = 7, eNpHR3.0 n = 8) and Control (eYFP n = 8; eNpHR3.0 n = 7). Block groups received Stage I training on Day 10 – 13 involving presentation of CSA (85dB 10Hz clicker) co-terminating with a 0.5 s, 0.6mA footshock. There were 4 presentations each day with an ITI between 600 – 720 s. The Control group received VI60 lever press training instead. Rats were tethered to dummy patch cables on the first day of Stage I (Day 10). All rats received Stage II training on Days 14 and 15. Prior to each session, rats were tethered to patch cables that outputted at least 10mW of 625 nm light. In Stage II, CSA was simultaneously presented with CSB (1Hz flashing LED) and this compound cue coterminated with a 0.5 s, 0.6mA footshock. There were 4 presentations each day with an ITI between 600 – 720 s. Photoinhibition was 1.5 s in duration, beginning 0.5 s prior to shock onset and terminating 0.5 s after shock offset, On Day 16, rats were tested. CSB was presented with an ITI of 900 s in a 70 min session.

### Immunohistochemistry

Localization of ChR2, eNpHR3., gCaMP7 or eYFP expression was verified using diaminobenzidine (DAB) immunohistochemistry. Free-floating brain tissue was washed in PB, pH 7.4 and rinsed in alcohol (50%), alcohol containing hydrogen peroxide (3%) and normal donkey serum (NDS; 5%) in PB, pH 7.4, for 30 min each. Sections were then incubated in rabbit anti-GFP (1:2000; Life Technologies) diluted in PBTX containing 2% NDS. for 24 h at room temperature. After washing off unbound primary antibody, sections were incubated overnight (at room temperature) in biotinylated donkey anti-rabbit IgG (1:5000; Jackson ImmunoResearch; Cat#711-065-152; RRID:AB_2540016) diluted in 2% NDS PBTX. After washing off unbound secondary antibody, sections were incubated for 2 h (at room temperature) in ABC reagent (Vector Elite Kit 6μ l/ml avidin and 6μl/ml biotin; Vector Laboratories). Brown GFP-IR cells were revealed using a DAB reaction, with peroxide being generated by glucose oxidase. In this DAB reaction, sections were washed in PB, pH 7.4, followed by 0.1 M acetate buffer, pH 6.0, and then incubated for 15 min in a 0.1 M acetate buffer, pH 6.0, solution containing 0.025% DAB, 0.04% ammonium chloride, and 0.02% D-glucose. The peroxidase reaction was started by adding 0.1μl/ml glucose oxidase and stopped using acetate buffer, pH 6.0. Brain sections were then washed in PB, pH 7.4, and mounted onto gelatin-treated slides, dehydrated, cleared with histolene, and cover slipped with Entellan (Proscitech, Kirwin, Australia). ChR2, eNpHR3.0, gCaMP7 and eYFP expression sites and cannula placements were determined under a microscope and plotted onto Illustrator (Adobe) templates using boundaries defined by Paxinos and Watson (2007). Rats with unilateral expression/cannula placements or expression/cannula placements outside the boundaries of the BLA were excluded from data analysis.

### Experimental design and Statistical Analyses

Suppression ratios were calculated as SR = a/(a+b) (Annau and Kamin, 1961), where a is the number of lever presses during the CS and b is the number of lever presses the minute prior to the CS (the pre-CS period). An SR of 0.5 indicates no suppression (equal number of lever presses during the CS and pre-CS period) whereas an SR of 0 indicates complete suppression, or asymptotic fear. These data and electrophysiology data were analysed via repeated measures ANOVA.

Fiber photometry signals were extracted and down sampled (15.89Hz) prior to further signal processing. The isosbestic signal was regressed onto the Ca2+-dependent signal to create a fitted isosbestic signal, and a fractional fluorescence signal ΔF/F was calculated via subtracting fitted 405 nm signal from 465 nm channels and then further dividing by the fitted 405 nm signal. High-pass 90 s) and low-pass filtering (3Hz) was conducted. ΔF/F signals around behavioral events (e.g. CS onset, US onset etc) were isolated and aggregated; 5 seconds before each event was used as baseline and the 7 s following each event was defined as the event transient. A bootstrapping confidence interval (CI) procedure (95% CI, 1000 bootstraps) was used to determine significant event-related transients within this window (Jean-Richard-dit-Bressel et al., 2020). A distribution of bootstrapped ΔF/F means was generated by randomly resampling from trial ΔF/F waveforms, with replacement, for the same number of trials. A confidence interval was obtained per timepoint using the 2.5 and 97.5 percentiles of the bootstrap distribution, which was then expanded by a factor of sqrt(n/(n-1)) to adjust for narrowness bias. Significant transients were defined as periods whose 95% CI did not contain 0 (baseline) for at least 0.5 secs. Areas under the curve for event transients were calculated by approximating the integral (trapezoidal method) of the isolated normalized ΔF/F curves. We analysed AUCs defined by subject means and by trial means via repeated measures ANOVA.

## Results

### AAV expression is specific to BLA PV neurons

We used immunohistochemistry to validate cell-type specificity of AAV expression. PV-Cre rats received BLA infusions of Cre-dependent eNpHR3.0-eYFP (n = 5) or Cre-dependent eYFP (n = 3) and two-colour immunofluoresence was used to process BLA sections for eYFP, PV, and eYFP+ PV immunoreactivity (-IR) (**Figure 1A**). There was robust and selective expression of these AAVs in BLA PV neurons, with some encroachment of the AAV into adjacent cortex. There was an average of 116.8 (SEM = 12.4) PV-IR neurons and 54.5 (SEM = 12.2) eYFP-IR neurons per animal with an average of 44% (SEM = 7) of BLA PV-IR neurons transduced. The majority (mean = 98%, SEM = 1) of eYFP-IR neurons also expressed PV-IR (mean dual-labelled neurons per animal= 53, SEM = 11.8) (**Figure 1C, D**). This confirms the utility of the PV-Cre rat to target BLA PV neurons.

**Figure 1.**
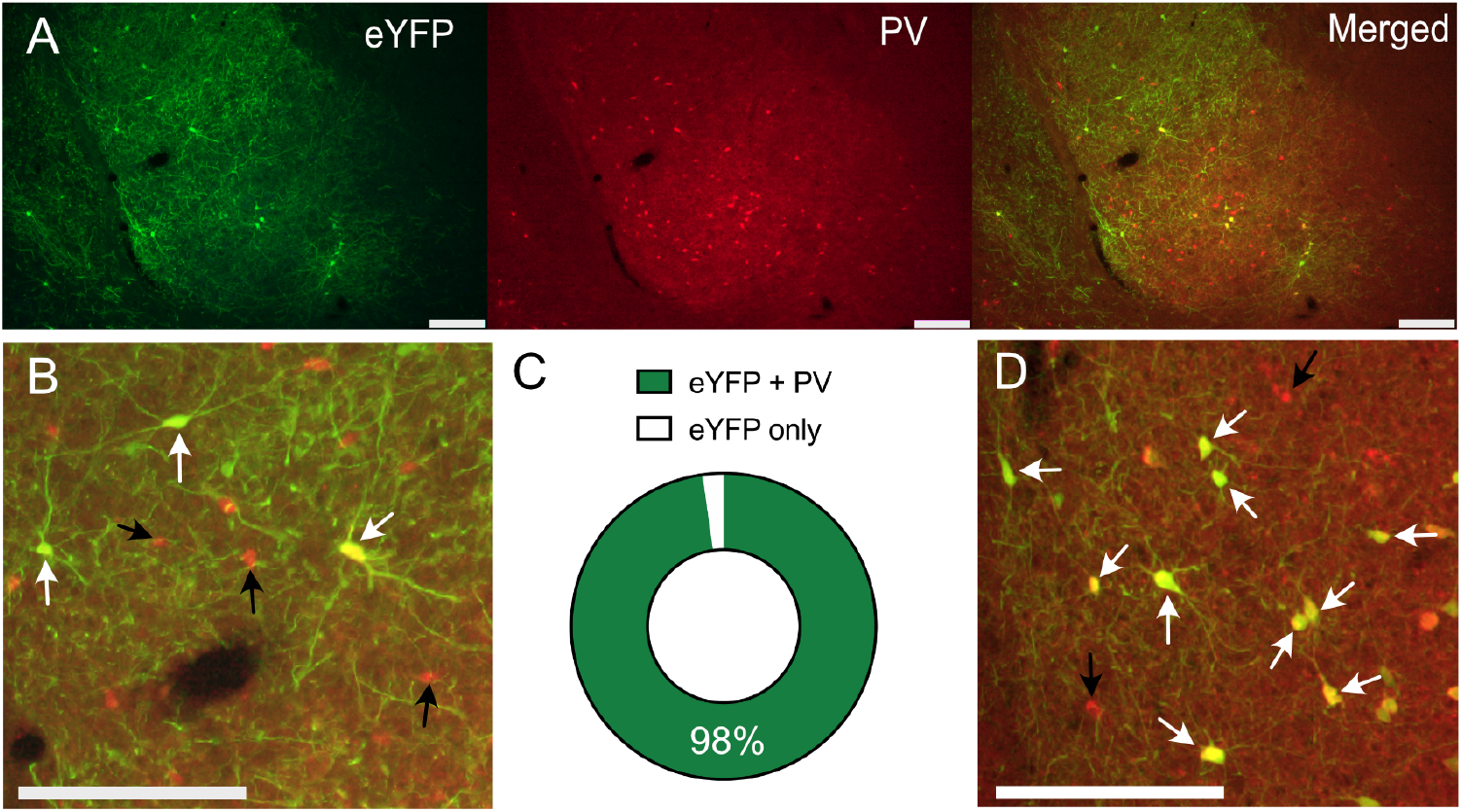
PV neuron specific expression. PV-Cre rats received BLA infusions of Cre-dependent eNpHR3.0-eYFP (n = 5) or Cre-dependent eYFP (n =3). **A**, Expression of eNpHR3.0-eYFP in BLA PV neurons. **B**, Higher magnification image showing dual labelled eNpHR3.0-eYFP/PV neurons. Dual labelled neurons shown with white arrows, single PV-labelled neurons with black arrows. **C**, The majority of eYFP neurons were co-labelled with PV. **D**, Higher magnification image showing dual labelled eYFP/PV neurons (white arrows) and single PV-labelled neurons with black arrows. Scale bars = 200 μm.

### Electrophysiological characterization of BLA PV neurons

Next, whole-cell patch-clamp recordings were made from eNpHR3.0-eYFP and ChR2-eYFP expressing BLA neurons from PV-Cre rats (**Figure 2A**). These neurons had membrane resistance (211 ± 15 MΩ), membrane time constant (18 ± 3 ms), brief APs (0.51 ± 0.04 ms) and prominent fast after hyperpolarizations (fAHPs) (16.0 ± 1.4 mV), with some firing action potentials in high frequency bursts with variable interburst intervals (‘stuttering’). These properties are consistent with the known properties of PV neurons (Rainnie et al., 2006; Woodruff and Sah, 2007b). Photoinhibition of eNpHR3.0-eYFP neurons evoked a rapid hyperpolarization persisting for the duration of the light and reliably suppressing PV neuron firing (**Figure 2C,D**). ChR2-eYFP expressing neurons were photostimulated (470 nm) with a sustained 1.5 s light pulse or a 25 Hz train of 20 ms light pulses for 1.5s (**Figure 2E**). Both protocols reliably evoked AP near the onset of the light pulse, but the 25 Hz train was better to sustain firing throughout the 1.5 s stimulation period (Two-way repeated measures ANOVA: time F (9, 45) = 5.472, p < 0.0001; light type F (1, 5) = 2.831, p = 0.1533; interaction F (9, 45) = 2.261, p = 0.0348; follow-up Sidak comparison p = 0.002) (**Figure 2E, 2G**)

**Figure 2.**
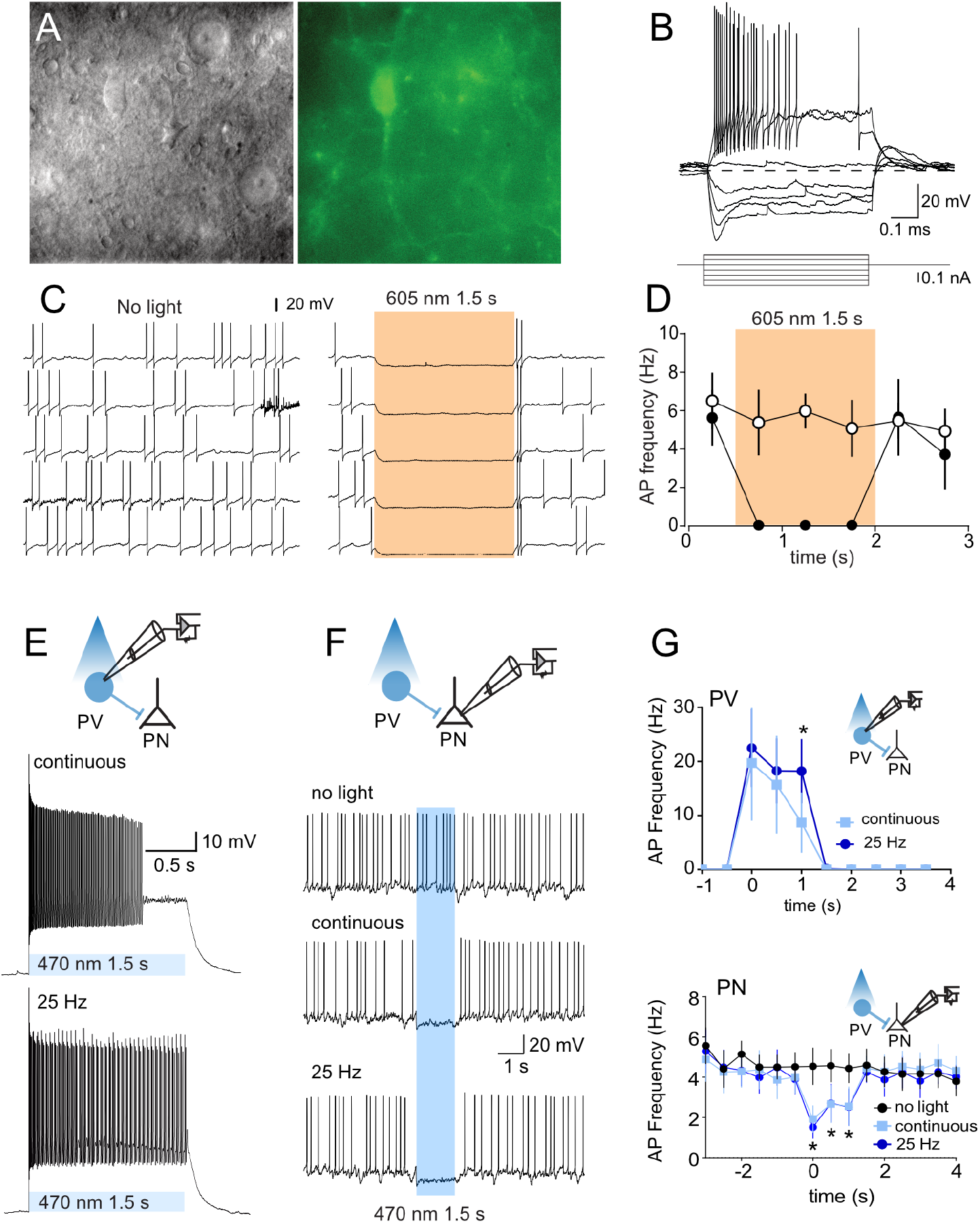
Light-evoked responses of eNpHR3.0 and ChR2 PV BLA neurons. **A**, Gradient contrast image (left) and green fluorescent image of PV neuron (right). **B**, Voltage response to current injections. **C,** Example of light-evoked suppression of PV neuronal firing. **D**, Mean (± SEM; n = 6) PV neuron firing frequency in presence and absence of light stimulation. Shaded areas indicate timing of light presentation. **E**, Example of light-evoked excitation of PV neuronal firing. **F**, Example of inhibition of BLA projection neuron (PN) firing by light-evoked excitation of PV neuron. **G**, Mean PV (n = 6) and PN (± SEM; n = 4) firing frequency in presence and absence of light stimulation. 25 Hz PV stimulation was better able to sustain PV neuron firing but there was no difference between these stimulation types in PN neuron inhibition. * p < .05.

We recorded from putative BLA projection neurons to assess the effect of PV neuron stimulation on BLA projection neuron firing (**Figure 2F, G**). These eYFP-negative neurons had a lower membrane resistance (116 ± 28 MΩ vs. 211 ± 15 MΩ; t = 3.0, p =0.009), broader APs (half-width 1.11 ± 0.06 ms vs 0.51 ± 0.04 ms; t = 7.792, p < 0.0001), and smaller fAHPs (half-width 6.7 ± 1.8 mV vs 16.0 ± 1.4 mV; t = 3.355, p = 0.0043), than the eYFP-positive PV neurons and their properties were consistent with BLA projection neurons (Sah et al., 2003). ChR2 stimulation of PV neurons with 470 nm light was sufficient to elicit IPSCs in projection neurons (data not shown) and to inhibit their firing (**Figure 2F, G**). We did not observe a difference in the effectiveness of continuous versus pulsed ChR2 stimulation in this inhibition of BLA projection neuronal firing (Two-way repeated measures ANOVA: time F (15, 45) = 4.464, p < 0.0001, light type F (2, 6) = 3.488, p = 0.0989; interaction F (30, 90) = 2.939, p = 0.0001).

### BLA PV activity at US delivery constrains fear learning

BLA PV interneurons exert strong perisomatic inhibition over BLA principal neurons. In mice, this inhibition constrains fear learning because optogenetic inhibition of PV neurons during the shock US augments, whereas optogenetic excitation during the shock US impairs, Pavlovian fear learning as measured by freezing (Wolff et al., 2014). We studied the effects of these same manipulations on Pavlovian fear learning in rats using conditioned suppression as the measure of fear. PV-Cre rats received AAVs encoding Cre-dependent ChR2, eNpHR3.0 or eYFP and fiber optic cannulae bilaterally into the BLA (**Figure 3A**). To establish a steady baseline of lever pressing for conditioned suppression, rats were initially trained to lever press for grain pellets for 10 days and then pre-exposed to an auditory stimulus for 2 days. They then received auditory fear conditioning via pairings of the auditory CS with an aversive shock US. We photoexcited or photoinhibited BLA PV neurons only during US delivery via a 465 (excite) or 625 (inhibit) nm light.

**Figure 3.**
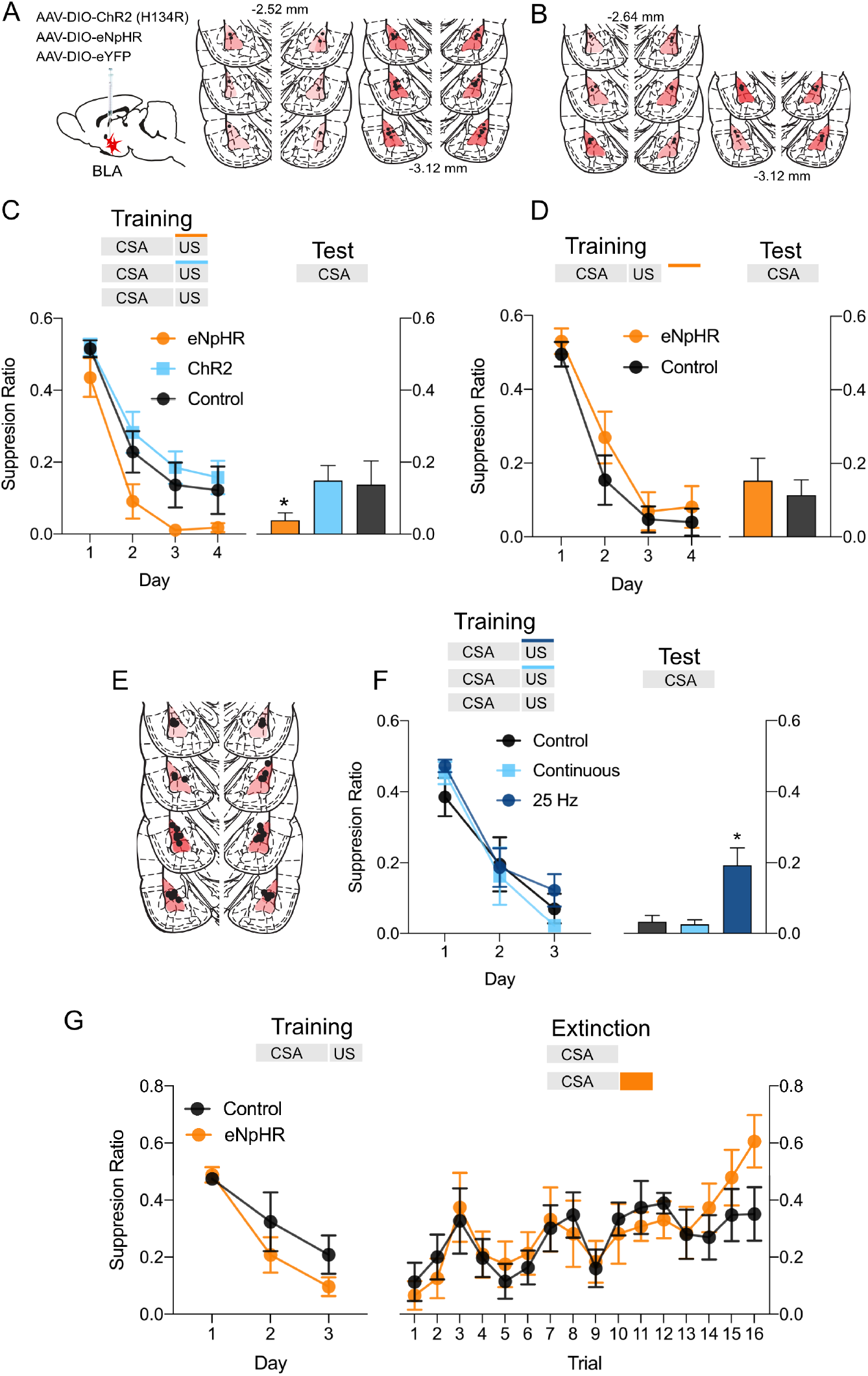
Bi-directional modulation of fear learning by optogenetic manipulation of BLA PV neurons. **A**, PV-Cre rats received BLA infusions of Cre-dependent eNpHR3.0 (n = 8) or ChR2 (n = 7) and fiber optic cannulae into the BLA. Controls were PV-Cre rats receiving cre-dependent eYFP (n = 4) or PV-Cre null rats receiving Cre-dependent ChR2 (n = 5) and fiber optic cannulae into the BLA. **B**, PV-Cre rats received BLA infusions of Cre-dependent eNpHR3.0 (n = 7) or eYFP (n = 8) and fiber optic cannulae into the BLA. **C**, BLA PV photoinhibition during shock US augmented fear learning. **D**, BLA PV photoinhibtion during the inter-trial interval had no effect. **E,F** 25Hz (n = 7) but not continuous (n = 9) BLA PV ChR2 photoexcitation during the shock US impaired fear learning compared to control (n = 9). **G**, Photoinhibition during shock omission had no effect on fear extinction. * p < 05.

Photoinhibition during the shock (eNpHR3.0) (n = 8) augmented fear learning compared to the control group (n = 9) (F(1, 21) = 5.70, p < 0.026), that did not interact with the main effect of day, F(1,21) = 0.052, p = 0.822 (**Figure 3C**). This augmented fear learning was preserved on test because the eNpHR3.0 group continued to show more fear when tested for long-term fear memory 24 hrs later (eNpHR3.0 vs ChR2: F(1,13) = 6.14, p = 0.028; vs Control F(1,15) = 1.85, p = 0.194). This augmentation of fear learning was specific to photoinhibition at the time of the shock US, because we conducted a separate experiment where rats received delivery of 625 nm light during the inter-trial interval rather than during the shock US **(Figure 3B**). There was no effect of this inter-trial interval photoinhibition on fear learning (F(1,13) = 1.27 p = 0.280) or long-term fear memory on test (F(1,13) = 0.79, p = 0.390) (**Figure 3D**).

BLA photoexcitation via ChR2 (n = 7), on the other hand, had no effect on fear learning (ChR2 vs controls: training: F(1,21) = 0.531, p = 0.474; test: F(1,21) = 0.001, p = 0.975) (**Figure 3C**). This was surprising because this same excitation impaired fear learning in mice (Wolff et al., 2014). Our electrophysiology had shown that a 25 Hz train of stimulation was better able to sustain PV neuron firing than continuous stimulation (**Figure 2E**), so it is possible that this lack of effect of PV ChR2 excitation on fear learning was due to use of continuous rather than pulsed stimulation. A separate experiment directly compared the effects of pulsed (25 Hz; n = 7) versus continuous (n = 9) ChR2 BLA PV neuron excitation during shock US delivery on fear learning (**Figure 3E, F**). Rats acquired fear to the auditory CS across fear conditioning (main effect of dat F(1,22) = 148.18, p < 0.001; no main effect of group F(2,22) = 0.275, p = 0.762; no group x day interaction: F(2,22) = 0.170, p = 0.844). On test, rats that had received 25 Hz ChR2 stimulation during US delivery showed significantly less long-term fear memory compared to continuous (F(1,22) = 5.38, p < 0.03) or no stimulation (F(1,22) = 14.07, p = 0.001). This shows that 25 Hz photoexcitation of PV neurons during the shock US impaired fear learning (**Figure 3F**). Consistent with our previous finding, there was no effect of continuous ChR2 stimulation on fear learning, F(1,22) = 2.35, p = 0.140 (**Figure 3F**). Taken together, these findings show bidirectional effects of BLA PV neuron photoinhibition and photoexcitation on fear learning.

### BLA PV neuron activity at US omission is not necessary for fear extinction learning

Having shown a role for BLA PV neurons at the time of shock US delivery in learning to fear, we next asked whether PV neurons are important at the time of shock omission for learning not to fear. Fear extinction learning is instructed by negative prediction errors at US omission (Rescorla and Wagner, 1972; Wagner and Rescorla, 1972). Fear extinction learning remodels PV perisomatic inhibitory synapses around BLA fear neurons (Davis et al., 2017; Trouche et al., 2013) and recruitment of these BLA PV neurons suppresses fear responding and activity in BLA fear neuronal ensembles during extinction retrieval (Davis et al., 2017; Ozawa et al., 2020). The activity of BLA PV neurons at the time of shock US omission is therefore an obvious candidate for instructing extinction learning, but whether PV neurons serve this role during US omission is unknown.

To study the role of BLA PV neurons in learning about shock omission during fear extinction, PV-Cre rats received AAVs encoding Cre-dependent eNpHR3.0 (n = 9) or eYFP (n = 7). They underwent auditory fear conditioning then extinction training (**Figure 3G**). BLA PV neurons were photoinhibited at the time of omission of the expected footshock US during extinction training. Across fear conditioning, rats acquired fear to the auditory CS (main effect of day, F(1,14) = 75.51, p < 0.001) with no differences between groups (no effect of group, F(1,14) = 1.45, p =0.248; or group x day interaction: F(1,14) = 2.80, p = 0.116) and they also extinguished this fear across extinction training (main effect of day: F (1,14) = 31.97, p < 0.001). Photoinhibition of BLA PV neurons at the time of shock omission had no effect this fear extinction learning (no main effect of group: F(1,14) = 0.025, p = 0.877; no group x day interaction: F(1,14) = 3.502, p = 0.082). So, in contrast to their role in fear learning at the time of US delivery, the activity of BLA PV neurons at the time of shock omission is not necessary for fear extinction learning.

### BLA PV neurons express the activity signatures of prediction error

BLA PV neurons are excited by the shock US to constrain activity in BLA projection neurons and constrain fear learning. We asked how the activity of BLA PV neurons changes across fear conditioning and across variations in aversive prediction error. To do this, we used fiber photometry (Gunaydin et al., 2014) to measure calcium transients in BLA PV neurons during an associative blocking task, the gold standard behavioral assessment of the role of prediction error in fear learning (Kamin, 1968; McNally and Westbrook, 2006). This allowed us to determine whether BLA PV neuron activity co-varies with US surprisingness in a task that precisely isolates the action of aversive prediction error.

PV-Cre rats received AAV encoding Cre-dependent gCaMP7s and fiber optic cannulae into the BLA (**Figure 4A, B**). There was selective expression of gCaMP7s in PV neurons (**Figure 4B**) with 28% of PV-IR neurons expressing gCaMP7. The associative blocking procedure uses a two-stage fear conditioning approach to isolate and assess prediction error. During Stage I, rats in the Block group received fear conditioning of an auditory CS, CSA. Rats in the Control group did not receive this training. This Stage I training establishes fear of CSA in group Block. Then, in Stage II, both groups received fear conditioning of a compound auditory (CSA) and visual (CSB) CS (CSAB). The Block group should not learn to fear CSB in Stage II because they have already learned that CSA signals shock, hence the shock US is expected in Stage II (i.e. prediction error is low). In contrast, the control group should learn to fear CSB during Stage II because the shock US is unexpected in Stage II (i.e. prediction error is high).

**Figure 4.**
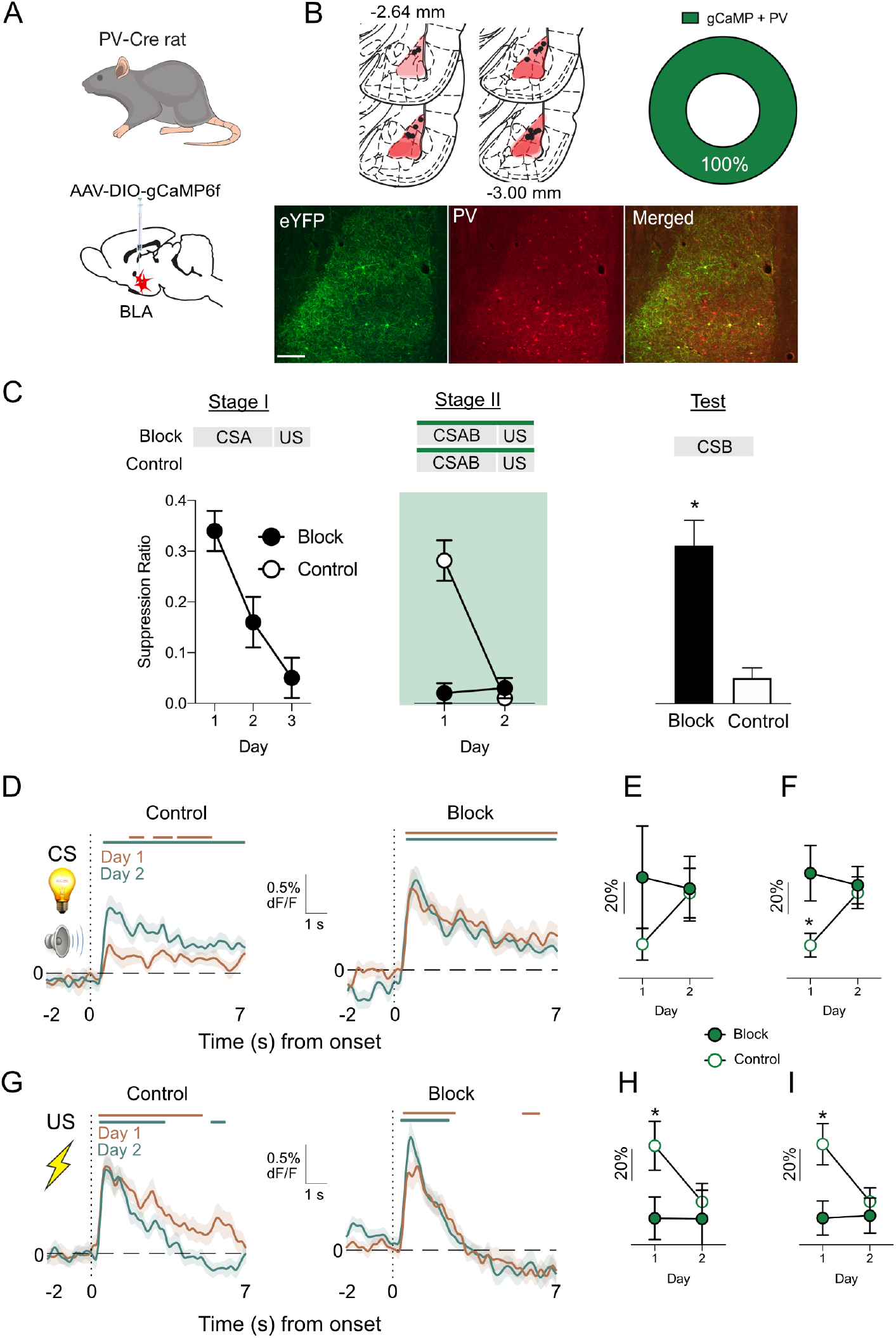
BLA PV neurons report aversive prediction errors. **A – B,** PV-Cre rats (n = 19) received AAV encoding Cre-dependent gCaMP7s and fiber optic cannulae into the BLA. Two-colour immunofluorescence showed gCaMP7s expression in PV neurons (n = 2). **C**, Blocking of associative fear learning. Group Block showed more fear to CSAB than Group Control (n = 9) in Stage II but Group Control learned to fear CSAB in Stage II. Group Control were more afraid of CSB on test than Group Block. **D,** Fibre photometry showing CS-evoked transients in BLA PV neurons during Stage II of associative blocking for groups Control and Block. Coloured bars above transients show periods significantly different to 0% ΔF/F (p <, .05). **E**, Subject-based AUC ΔF/F 0 – 7s after CS onset. **F**, Trial-based AUC ΔF/F 0 – 7s after CS onset. **G,** US-evoked transients in BLA PV neurons during Stage II of associative blocking for groups Control and Block. **H**, Subject-based ΔF/F 0 – 7s after US onset. **I**, Trial-based ΔF/F 0 - 7s after US onset. * p < .05.

Precisely this pattern of results was observed (**Figure 4C**). Group Block (n = 10) learned to fear CSA in Stage I (F(1,9) = 41.00, p < 0.001). Group Block also showed more fear to CSAB than Group Control (n = 9) in Stage II (main effect of group: F (1,17) = 41.56, p < 0.001), but Group Control did learn to fear CSAB in Stage II (main effect of day: F (1,17) = 25.56, p < 0.001; and group x day interaction: F(1,17) = 28.32, p < 0.001). Critically, on test, Group Control were more afraid of CSB than Group Block (F(1,17) = 18.67, p < 0.001). This shows the associative blocking of fear learning: the same aversive footshock US supported different amounts of fear learning, depending on whether it is expected or unexpected. Rats learned more fear to an unexpected (Group Control) than expected (Group Block) aversive US.

BLA PV neurons showed significant Ca^2+^ transients at both CS onset and US onset during Stage II fear conditioning (**Figure 4D,G**). Critically, these transients changed across Stage II. We measured area under the ΔF/F curve and found a significant three-way interaction between stimulus type (CS v US), day (Day 1 v Day 2) and group (Block v Control) for AUCs (trials: F (1, 60) = 12.70, p = 0.001; subjects F (1, 16) = 13.11, p = 0.002). Further analyses showed that US-evoked Ca^2+^ transients decreased across Stage II (main effect of day: trials, F(1, 62) = 5.197, p <0.026; subjects, F(1, 16) = 6.835, p < 0.019) and this decrease was greatest in group Control (day x group interaction: trials, F(1, 62) = 4.165, p < 0.046; subjects, F(1,16) = 6.616, p < 0.020) (**Figure 4H,I**). This shows diminution of US-evoked activity for group Control. Indeed, Group Control had significantly greater US-evoked transients than group Block on Day 1 (trials: F (1, 69) = 7.74, p = 0.007; subjects, F (1, 17) = 4.75, p = 0.04) but not Day 2 (trials, F (1, 66) = 0.32, p = 0.75; subjects, F (1, 16) = 0.21, p = 0.65) (**Figure 4H,I**).

CS-evoked activity tended to show the opposite pattern, with strongest changes detected in the trial-based analysis. The increase in CS-evoked transients was greater for group Control than for group Block (day x group interaction: trials, F(1, 63) = 7.9, p < 0.007; subjects, F(1,16) = 3.456, p = 0.082). Group Block had significantly greater CS-evoked transients than group Control on Day 1 (trials, F(1, 70) = 5.50, p = 0.022; subjects, F (1, 17) = 1.40, p = 0.25) but not Day 2 (trials, F (1, 66) = 0.10, p = 0.76; subjects, F (1, 16) = 0.01, p = 0.92) (**Figure 4E,F**).

These findings show US- and CS-evoked activity in BLA PV neurons during fear conditioning and that this activity scales with both prediction (i.e. CS-elicited activity) and prediction error (i.e. US-elicited activity). PV neurons were strongly excited by an unexpected US (high prediction error) and these responses decreased as the US became expected (low prediction error). At the same time, CS-evoked PV neuron responses were low when the CS was a poor predictor of the shock US and these responses increased across the course of conditioning as the CS was learned as a good predictor of shock US. These opposing changes in PV neuron responsiveness to the US and CS across the course of conditioning are the activity signatures of aversive prediction error (Herry and Johansen, 2014; McNally et al., 2011).

### Role of BLA PV neurons constrain learning across variations in prediction error

Having demonstrated the aversive prediction error sensitivity of BLA PV neurons, we next asked how BLA PV neurons control fear learning across these variations in prediction error. To do this, we photoinhibited BLA PV neurons at the time of the expected shock US during Stage II of the associative blocking procedure. PV-Cre rats received AAV encoding Cre-dependent eNpHR3.0 or eYFP and fiber optic cannulae into the BLA (**Figure 5A, B**). They then received the two-stage associative blocking procedure. BLA PV neurons were silenced during US delivery in Stage II. There were 4 groups: eNpHR3.0-Block; eYFP-Block; eNpHR3.0-Control; eYFP-Control.

**Figure 5.**
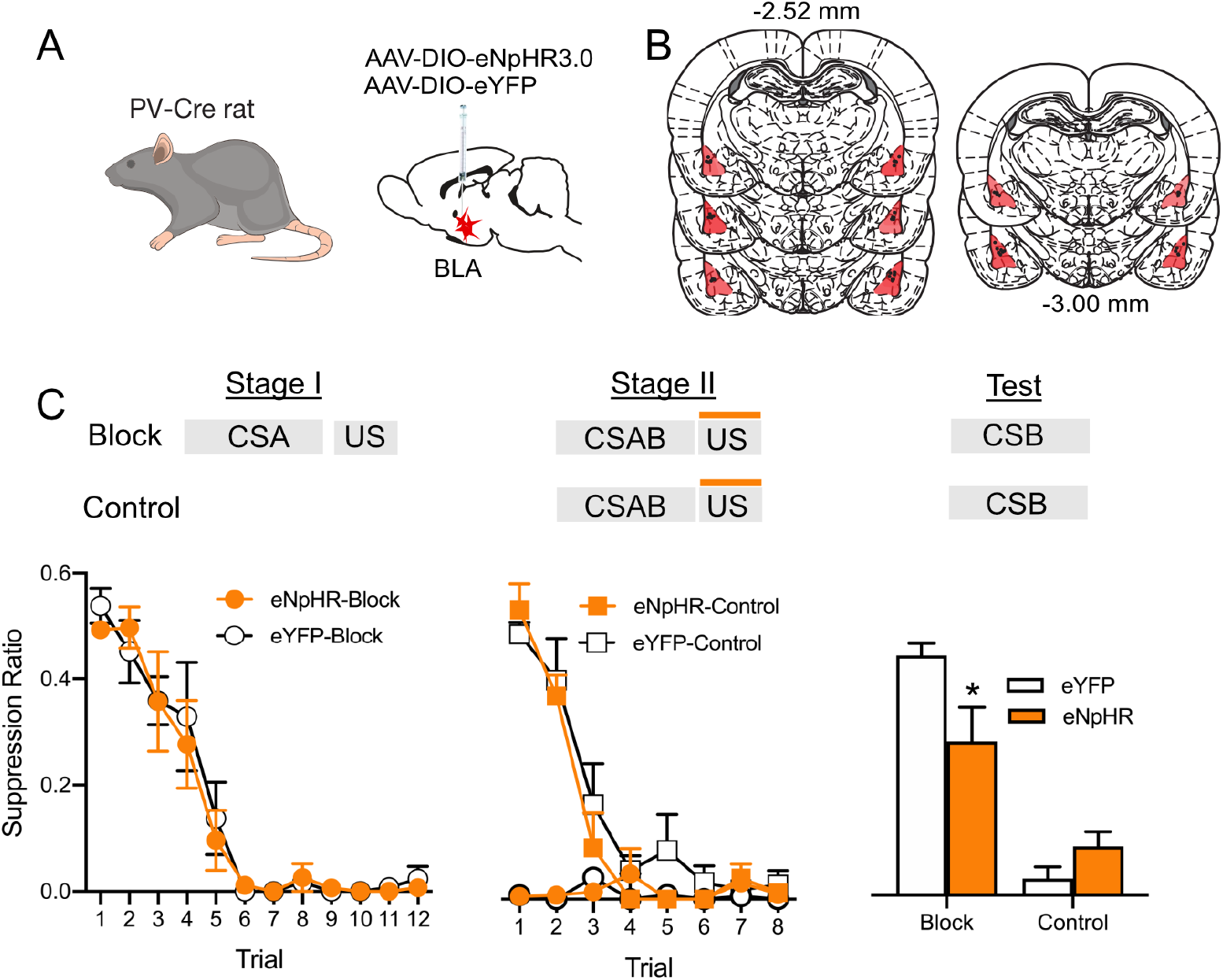
BLA PV neurons use aversive prediction error to constrain association formation. **A – B,** PV-Cre rats received BLA infusions of Cre-dependent eNpHR3.0 or eYFP and fiber optic cannulae into the BLA. **C,** eNpHR3.0-Block (n = 8) and eYFP-Block (n = 7) both learned fear to CSA in Stage I and eYFP-Control (n = 8) learned to fear CSAB during Stage II. Blocking was demonstrated at test. BLA photoinhibition during the Stage II shock US reduced blocking and restored fear learning to CSB in group eNpHR3.0-Block. * p < .05.

As expected, the Block groups acquired fear to CSA in Stage I (main effect of day: F(1,13) = 274.60, p < 0.001) (**Figure 5C**). There was no manipulation during Stage I and the Block-eYFP and Block-eNpHR3.0 groups did not differ during this stage (no main effect of group: F(1,13) = 0.069, p = 0.797; no group x day interaction: F(1,13) = 0.034, p = 0.857). During Stage II, the Block groups showed high levels of fear to CSAB whereas the Control groups learned to fear CSAB (main effect of group: F(1, 26) = 90.41, p < 0.05; main effect of day: F(1,26) = 49.94, p < 0.001; group x day interaction: F(1,26) = 43.23, p < 0.001) (**Figure 5C**). There was no effect of photoinhibition of BLA PV neurons during Stage II. At test, the Block groups showed less fear compared to Control groups (main effect group: F(1,26) = 65.42, p < .0001) – again demonstrating successful blocking and the powerful role of prediction error in constraining fear association formation. Importantly, silencing BLA PV neurons during the expected shock in Stage II reduced this blocking (group x virus interaction (F 1,26) = 8.51, p < 0.007; eNpHR3.0 – Block vs eYFP-Block F(1,13) = 9.06, p = 0.01) (**Figure 5C**). Thus, BLA PV neurons constrained fear learning during states of low prediction error (blocking). PV neuron inhibition did not augment learning in the control group in this experiment, in contrast to our earlier experiment. This is unsurprising. The experimental parameters, including the amount of training, US intensity, use of a compound CS, and conditions of testing were optimised to detect associative blocking not augmentation of *de novo* fear learning.

## Discussion

Here we used the PV-Cre rat (Wright et al., in preparation) to study the role of BLA PV neurons in Pavlovian fear conditioning and aversive prediction errors. We first confirmed the validity of the PV-Cre rat line. Cre-dependent AAVs robustly expressed in BLA neurons with high levels of selectivity to PV neurons. Although it is worth noting that less than half of the BLA PV neurons we identified expressed the AAV constructs. This relatively low expression was observed across two different AAVs, so does not appear to be specific to choice of AAV, but whether it is unique to the PV-Cre rat or whether it also observed in the PV-Cre mouse is unknown. Regardless, this may contribute to modest behavioral effects of PV manipulations here and elsewhere (Wolff et al., 2014). We confirmed the electrophysiological properties of these PV neurons as well as their ability to inhibit action potential firing in BLA projection neurons. We show a role for US-evoked activation of BLA PV neurons in constraining Pavlovian fear association formation. We also extend this role to show that BLA PV neuron activity at US omission is not necessary for fear extinction learning. Together, these findings confirm the utility of the PV-Cre rat for studying BLA PV neuron contributions to fear learning. These results also support the claim that BLA PV neurons constrain shock US driven fear learning (Wolff et al., 2014).

However, it is important to recognise that PV neurons are highly heterogeneous. PV neurons have diverse firing properties and form multiple networks in the BLA, including PV neuron → projection neuron, projection neuron → PV neuron, and PV neuron → other interneuron networks (Woodruff and Sah, 2007b). There are also differences in fear conditioning-related plasticity between PV neurons located across the amygdala (Lucas et al., 2016). This considerable diversity in how PV neurons contribute to the intrinsic circuitry of the BLA, and in their plasticity, likely allow them to perform a variety of functions in fear learning and amygdala beyond those studied here (Morrison et al., 2016).

There was pronounced CS- and US-evoked activity in rat BLA PV neurons across fear conditioning. CS-evoked activity in BLA PV neurons is consistent with past reports (Krabbe et al., 2019; Wolff et al., 2014) but past findings regarding US-evoked activity of BLA PV neurons during fear conditioning are mixed. PV neurons show heterogenous responses to noxious stimuli (Bienvenu et al., 2012). In fear conditioning, single-unit recordings show that some PV neurons can be inhibited by a footshock US to disinhibit BLA projection neurons allowing fear association formation (Wolff et al., 2014). By contrast, in calcium imaging many PV neurons are excited by footshock (Krabbe et al., 2019) and this is consistent with findings that footshock causes rapid inhibitory synaptic input to restrict action potential generation and firing frequency of BLA projection neurons (Windels et al., 2010). We show that PV neurons are robustly activated by the shock US, that *in vitro* excitation of these neurons inhibited BLA projection neurons, and that silencing PV neurons during shock presentation but not shock omission affected fear learning. Although fibre photometry did not allow us to resolve the activity of individual BLA PV neurons, our findings do show a key role for shock-US evoked PV neuron activity in controlling fear learning.

The activity of BLA PV neurons during Pavlovian fear conditioning reports aversive prediction error. PV neurons were strongly excited by an unexpected US and these responses decreased as the US became expected. At the same time in the same animals, CS-evoked PV neuron activity increased across the course of conditioning and was greatest when the CS was a better predictor of the shock US. These opposing changes in PV neuron responsiveness to the US and CS across the course of conditioning are the hallmarks of prediction error. This prediction error may be critical for instructing learning-related plasticity in PV neurons (Lucas et al., 2016).

Nonetheless, our findings argue strongly against the possibility that BLA PV neurons are themselves involved in the computation of this aversive prediction error. The key model preparation for understanding neural prediction errors is dopamine neurons in the VTA. During appetitive conditioning, VTA dopamine neurons are sensitive to reward prediction errors. They are robustly recruited by surprising or unexpected rewards but not by expected rewards (Schultz, 2006; Schultz et al., 1997). At the same time, CS-evoked activity in dopamine neurons increases across the course of conditioning and is greatest when the CS is a good predictor of reward. This prediction error sensitivity is achieved, at least in part, by VTA GABA neurons that gate activity in VTA dopamine neurons. VTA GABA neurons show a profile of ramping activity at CS onset, peaking at the moments of US delivery to inhibit both US-evoked responses in VTA dopamine neurons and reward learning (Cohen et al., 2012; Eshel et al., 2015). Despite their significant inhibitory synaptic inputs to BLA projection neurons and their sensitivity to prediction error, our data suggest that it is unlikely that BLA PV neurons similarly gate BLA projection neurons to compute an aversive prediction error. CS evoked responses in PV neurons peaked at CS onset and declined thereafter. Moreover, if the activity of BLA projection neurons is gated by PV neurons to compute a prediction-error corrected US signal, then we expect projection neurons to require the greatest inhibition and PV neurons to show the greatest excitation at the time of the expected shock US. Again, we did not observe this. Instead, we found the opposite. US-evoked activity in PV neurons was greatest for unexpected shock USs and decreased as prediction error decreased. Thus, we conclude it is unlikely that BLA PV neurons determine aversive prediction error sensitivity in the BLA.

Instead, our results suggest that reporting of aversive prediction errors is a common feature of otherwise distinct BLA cell-types. The same profile of prediction-error related changes that we observed in PV neurons have been reported in rat single unit recordings of slow (putative projection) and fast (putative interneurons) firing BLA neurons (Johansen et al., 2010), in mouse single-unit recordings of identified VIP interneurons (Krabbe et al., 2019), and in mouse BLA (McHugh et al., 2014) as well as human amygdala fMRI BOLD signals (Eippert et al., 2012; Michely et al., 2020). So, aversive prediction error sensitivity is not a specific or unique feature of individual BLA cell types. Rather, it appears to be a common feature of diverse BLA cell-types, despite these cell types having distinct roles in fear learning (Letzkus et al., 2015; Tovote et al., 2015).

This preservation of prediction error sensitivity across diverse BLA cell types suggests that the computation of aversive prediction error during fear conditioning is extrinsic to the BLA. The most plausible possibility is that diminution of US-evoked activity as the US becomes expected is linked to circuits involving the midbrain periaqueductal gray (Cole and McNally, 2007, 2008; Fanselow, 1998; McNally, 2005; McNally and Cole, 2006; McNally et al., 2011; McNally and Westbrook, 2006; Ozawa et al., 2017; Walker et al., 2020; Wright and McDannald, 2019) and that a prediction error signal is then relayed to and distributed across different BLA cell-types via the midline thalamus and prefrontal cortex (Dunsmoor et al., 2008; Eippert et al., 2012; Furlong et al., 2010; Sengupta and McNally, 2014). A different possibility is that PV neurons may be sensitive to prediction error via their receipt of direct excitatory inputs from BLA projection neurons (Woodruff and Sah, 2007a, b) which have also been shown to report aversive prediction errors (Johansen et al., 2010; Ozawa et al., 2017).

Regardless, different populations of BLA neurons use an aversive prediction error signal in a cell-type specific manner to instruct and regulate fear association formation. BLA projection neurons (Johansen et al., 2010; Ozawa et al., 2017) and VIP interneurons (Krabbe et al., 2019) use this prediction error signal to form and store fear associations. In marked contrast, PV interneurons use this signal to constrain fear association formation.

## Acknowledgments

Data reported here are archived in the UNSW Long Term Data Archive (ID:). This work was supported by the Australian Research Council (DP170100075) and UNSW School of Psychology. We thank Pankaj Sah for helpful comments on this manuscript.

## Author contributions

### Conceptualization

JOY, JMP, GPM.

### Experimentation

JOY, CC, JMP.

### Formal Analysis

JOY, CC, JMP.

### Writing - original draft

JOY, GPM.

### Writing - Review & Editing

All authors.

### Funding acquisition

GPM, JMP.

## Notes

### Competing Interest Statement

The authors have declared no competing interest.

